# Growth assessment of juvenile oil palm (*Elaeis guineensis* Jacq.) intercropped with fruit vegetables in a rainforest zone of Nigeria

**DOI:** 10.1101/2020.05.31.126714

**Authors:** Ayodele Samuel Oluwatobi

**Affiliations:** Department of Biological Sciences, Crown-Hill University, Eiyenkorin, Ilorin, Nigeria

**Keywords:** crop association, development, immature, performance, spacing, vegetables, weed

## Abstract

Many farmers spend a lot of money in the control of weeds during the long juvenile phase of oil palm. Intercropping is a popular practice employed by farmers to increase productivity of wide alley and overcome weed problem. However, improper intercropping of *Elaeis guineensis* with other crops has impaired the growth and development of the oil palm due to competition for environmental resources. The study was conducted to investigate the impact of intercropping on the growth of juvenile oil palm for 2 years. The research commenced during the rainy season of 2016 at an established juvenile oil palm plantation in Ala, Akure-North Local Government of Ondo State. Four fruit vegetables were intercropped separately within the alley of the plantation at 1, 2 or 3 m away from the oil palms in a randomized complete block design. Growth parameters of intercropped and non-intercropped juvenile oils were compared. Results of the study revealed that at 16 weeks after intercropping (WAI), the intercropped oil palm recorded better growth performance with higher values of canopy spread, number of frond, number of leaflets and trunk height (218.20, 37.00, 87.48 and 38.17 cm) respectively, than the sole oil palms (214.67, 32.83, 72.89 and 31.67) respectively. There was no significant difference in all the growth parameter examined except canopy height at (P<0.05) level of significance. Juvenile oil palm cultivated in rainforest agroecological zone of Nigeria can be intercropped with fruit vegetables without any deleterious effect when intercropped at minimum of 1 m away from the oil palms.

## 1. Introduction

Growing a number of other food crops in association with juvenile oil palm trees is a widespread practice in most oil palm growing areas in the country. Oil palm is a wide spaced perennial crop with a long juvenile period of 3-5 years depending on cultivars (Igene *et al.*, 2015). The space in the inter- and intra-rows can be put to use to create income during the long juvenile period of the crop. The spacing habit and growth pattern (9 × 9 m triangular spacing) of oil palm plantation permit a variety of annual and perennial crops to be cultivated with and under its canopy during the young stage of the oil palm (NIFOR, 2008). The benefits of intercropping with oil palms in the field according to NIFOR (2008) are adding value to the oil palm when the food crops are harvested and sold, and particularly so in the early years when the oil palm has not started to yield fruit bunches. Ibeawuchi (2007) reported that intercropping suppresses weeds and gives yield advantage and stable yield overtime. He opined that when suitable crops are grown in proximity, it promotes positive interaction among them.

Secondly, the planting of the food crops between inter-rows of the oil palm facilitates field maintenance of the plantations, resulting in reduction in overall maintenance cost. Hence, the double cost of maintenance is avoided (Igene *et al.*, 2015).

Oil palm can successfully be intercropped with food crops. Nuertey *et al.* (2000) indicated that it is profitable to intercrop oil palm with food crops especially for the first three to four years when the palms are not fruiting as compared to sole cropping. Farmers are able to get enough money from the intercrop to sustain their family and also to maintain the farm.

The relative advantage of intercropping oil palm with food crops, suggests that intercropping systems may be most suitable for small-scale producers with limited resources to purchase large land to develop oil palm and food crops separately (Okyere *et al.*, 2014). Intercropping is also important because it helps smallholder (SH) farmers who face labour constraints as they have the potential to reduce weeding frequency (Mashingaidze *et al*., 2000).

Nwawe *et al.* (2014) studied the economic analysis of economic analysis of oil palm and food crop enterprises in Edo and Delta State Nigeria; they stated that sustainable and stable mixed oil palm food crop enterprise in Nigeria requires that farmers are guided by for the choice of oil palm food crop combination.

Udosen *et al.* (2006) investigated the performance of oil palm under food crop combinations in four-year old oil palm in derived savannah zone of Nigeria. They asserted that there is benefit in appropriate cropping mixture with immature oil palm.

However, intercropping of juvenile oil palm with these food crops result in competition between the juvenile oil palm and the food crops for resources such as water, space and nutrients. This competition could result in adverse effect if there are limited resources in the environment. It is therefore important to assess the implication of the intercrop on the growth of the juvenile oil palm during the long juvenile phase of the crop. Therefore, the objective of this study is to evaluate the effects of intercropping on the growth of juvenile oil palm in rainforest agroecological zone of Nigeria. Hence, farmers can be educated on the proper way to carryout intercropping with juvenile oil palm in a way that would not cause deleterious effect on their growth.

## 2. Materials and methods

Description of study area: The field experiment was conducted within an established juvenile oil palm plantation located at Ala in Akure-North Local Government Area of Ondo State. The oil palm plantation is located at a coordinate range of Latitude 7.093° N, Longitude 5.354°E (N7°5’ 35.59837” E5°21’ 15.47179”), and Latitude 7.09302 Longitude 5.35422 (N7°5’ 34.8857” E5°21’ 15.19177”) in the tropical rain forest region of Nigeria. It has two distinct seasons namely: dry and rainy seasons. Rainy season is between April and November and dry season is between November and March. Annual rainfall varies from 1150 to 2550 mm. Temperature is moderately high year-round and range between 22°C and 34°C with daily average of 30°C (Ogunrayi *et al.*, 2016). Top soil (0-15 cm) was collected for soil test before and after intercropping to establish the physicochemical properties of the soil. These properties include: soil pH, total carbon, organic matter, electrical conductivity, exchangeable cations (Na^+^, Mg^2+^, K^+^, Ca^2+^ and titratable acidity or acid value), nitrate content, phosphorus content (Mussa *et al*, 2009), particle size (clay, sand and silt) according to the method of Kettler *et al.*(2001), and soil type (Olabisi *et al.*, 2009). Electrical conductivity, total carbon and total organic matter were determined according to the methods of Wagh *et al.* 2013. Exchangeable cations, titratable acidity, nitrate and phosphate contents were determined according to the methods of Reeuwijk (2002), Czinkota *et al.* (2002), Ahmed (2009) and Mussa *et al.* (2009), respectively. Soil particle size was determined according to the method of Kettler *et al.* (2001).

The juvenile oil palm trees in the plantation were planted at a plant spacing of 6m × 6 m. Four fruit vegetables: (i) tomato (accessions NGB 01665 and NG/AA/SEP/09/053); (ii) pepper (NGB 01312 and NGB 01641); (iii) okra (NGB 01184); and (iv) eggplant (NGB 01737) were obtained at the National Centre for Genetic Resources and Biotechnology (NACGRAB) research institute, Ibadan, Oyo State and intercropped within alley of juvenile oil palms (240 m^2^). The different accessions of the fruit vegetables were intercropped at 1, 2 or 3 m away from the juvenile oil palm at spacing of 1 × 1 m within the alley, in a randomized complete block design with four replicates each. The blocks were represented by the replicates and the treatments (intercropping distances) were assigned once within each block of the fruit vegetables-juvenile oil palm intercrop plots. The control plot was without juvenile oil palm.

The field experiment was carried when the juvenile oil palms were 2 years old.

Measurements were taken from sole and intercropped juvenile oil palms every 4 weeks (4 weeks, 8 weeks and 16 weeks after intercropping).

Data collection and processing:

Growth variables collected on the juvenile oil palm include:

Number of fronds: This was determined by counting the number of fronds on each juvenile oil palm tree.

Average length of fronds: This was obtained by measuring with a meter rule the length of the fronds and finding the average.

Average number of leaflets: This was obtained by counting the number of leaflets on fronds and finding the average.

Average canopy spread determination: Canopy spread of the immature oil palms was measured according to Spoke Method (Blozan, 2004) with the use of graduated coloured plank or wooden rod (tar rod). Ten measurements were taken from the midpoint of the trunk to the outer extremities of the crown. These were averaged to get the result of the average canopy or crown spread. Canopy spread measurements were taken every 4 weeks (4, 8, 12 weeks and until final harvesting).

Trunk height: This was measured with the use of graduated coloured plank (tar rod) (Blozan, 2004).

Canopy height: It was also measured with the use of graduated coloured plank (tar rod) (Blozan, 2004).

Statistical analysis: T-test was used to analyze data gathered from the study using Statistical Package for Social Sciences (SPSS: version 17.0). Graphs were plotted using Origin (version 7.0) software.

## Results

The result on physical and chemical properties of the soil is shown in Table 1a-b.

**Table 1a:** Physical and chemical properties of the experimental plot before planting and after harvesting

| Properties |  |  |  |  |  |  | Particle size distribution |  |  |
| --- | --- | --- | --- | --- | --- | --- | --- | --- | --- |
| Plot |  | pH value | Total organic | Organic matter (%) | Nitrate ion (mg/l) | Available phosphorus (%) | Sand (%) | Silt (%) | Clay (%) |
| Pre-planting | NGB 01665 | 6.02 | 1.47 | 2.542 | 1.44 | 1.84 | 47.60 | 12.48 | 39.92 |
|  | NG/AA/SEP/09/053 | 5.10 | 1.40 | 2.421 | 1.27 | 1.65 | 58.45 | 13.64 | 27.91 |
|  | NGB 01312 | 5.86 | 1.46 | 2.524 | 1.41 | 1.78 | 47.36 | 14.21 | 38.43 |
|  | NGB 01641 | 5.99 | 1.41 | 2.438 | 1.40 | 1.70 | 59.32 | 13.84 | 26.84 |
|  | NGB 01184 | 6.08 | 1.35 | 2.334 | 1.23 | 1.62 | 46.76 | 12.78 | 40.46 |
|  | NGB 01737 | 6.09 | 1.40 | 2.421 | 1.28 | 1.68 | 48.42 | 13.76 | 37.82 |
|  | Control (sole) | 6.36 | 1.58 | 2.732 | 1.65 | 1.88 | 44.68 | 14.35 | 40.97 |
| Post-harvest | NGB 01665 | 6.38 | 1.49 | 2.576 | 1.45 | 1.96 | 49.62 | 13.88 | 36.50 |
|  | NG/AA/SEP/09/053 | 6.23 | 1.36 | 2.351 | 1.25 | 1.66 | 50.84 | 13.90 | 35.26 |
|  | NGB 01312 | 2.04 | 2.64 | 4.565 | 2.78 | 2.98 | 48.66 | 13.69 | 37.65 |
|  | NGB 01641 | 4.86 | 1.34 | 2.317 | 1.23 | 1.60 | 43.98 | 14.62 | 41.40 |
|  | NGB 01184 | 5.62 | 1.40 | 2.421 | 1.27 | 1.67 | 45.86 | 13.86 | 40.28 |
|  | NGB 01737 | 6.02 | 1.41 | 2.438 | 1.40 | 1.72 | 47.73 | 14.36 | 37.91 |
|  | Control (sole) | 6.23 | 1.50 | 2.594 | 1.46 | 1.97 | 42.87 | 14.38 | 42.75 |

**Table 2b:** Physical and chemical properties of the experimental plot before planting and after harvesting

| Properties |  | Exchangeable cations |  |  |  |
| --- | --- | --- | --- | --- | --- |
| Plot |  | Ca<sup>2+</sup>(mg/kg) | Mg<sup>2+</sup> (mg/kg) | Na<sup>+</sup> (mg/kg) | K<sup>+</sup> (mg/kg) |
| Pre-planting | NGB 01665 | 1.41 | 0.19 | 0.84 | 0.191 |
|  | NG/AA/SEP/09/053 | 1.39 | 0.19 | 0.82 | 0.188 |
|  | NGB 01312 | 1.40 | 0.18 | 0.83 | 0.190 |
|  | NGB 01641 | 1.40 | 0.18 | 0.82 | 0.192 |
|  | NGB 01184 | 1.39 | 0.20 | 0.80 | 0.186 |
|  | NGB 01737 | 1.37 | 0.17 | 0.80 | 0.185 |
|  | Control (sole) | 1.53 | 0.22 | 0.87 | 0.194 |
| Post-planting | NGB 01665 | 1.42 | 0.20 | 0.85 | 0.192 |
|  | NG/AA/SEP/09/053 | 1.35 | 0.16 | 0.78 | 0.188 |
|  | NGB 01312 | 1.58 | 0.26 | 0.89 | 0.198 |
|  | NGB 01641 | 1.33 | 0.14 | 0.76 | 0.185 |
|  | NGB 01184 | 1.38 | 0.18 | 0.81 | 0.187 |
|  | NGB 01737 | 1.39 | 0.19 | 0.82 | 0.186 |
|  | Control (sole) | 1.43 | 0.20 | 0.85 | 0.191 |

The results on the effects of intercropping fruit vegetables with juvenile oil palm during second year of plantation establishment are given in the Figure 1-6.

**Fig. 1:**
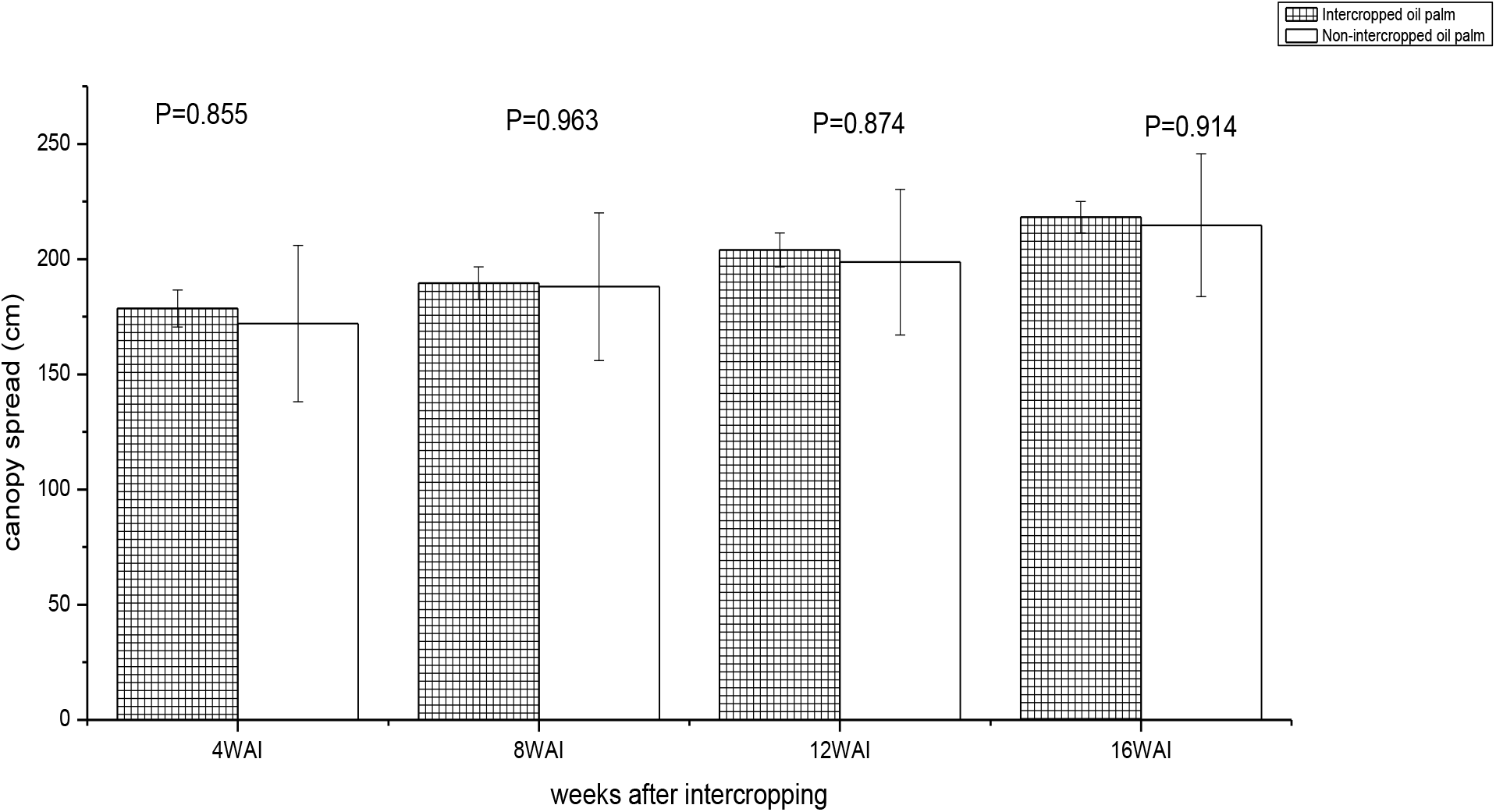
Canopy Spread of intercropped and sole juvenile oil palms at 4, 8, 12 and 16 weeks after intercropping. (n= 6) (2 years old).

**Fig. 2:**
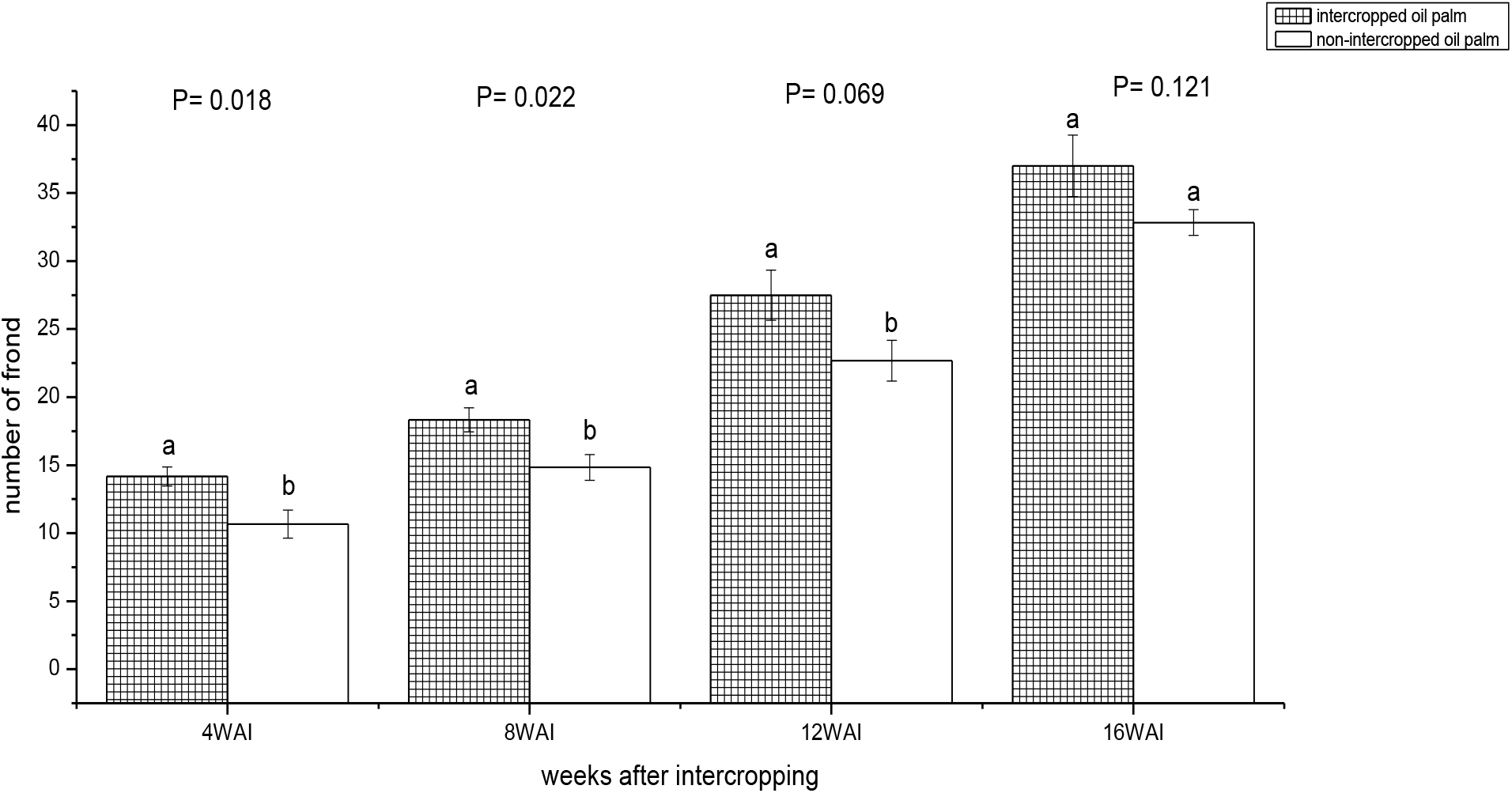
Number of fronds of intercropped and sole juvenile oil palms at 4, 8, 12 and 16 weeks after intercropping. (n= 6) (2 years old).

**Fig. 3:**
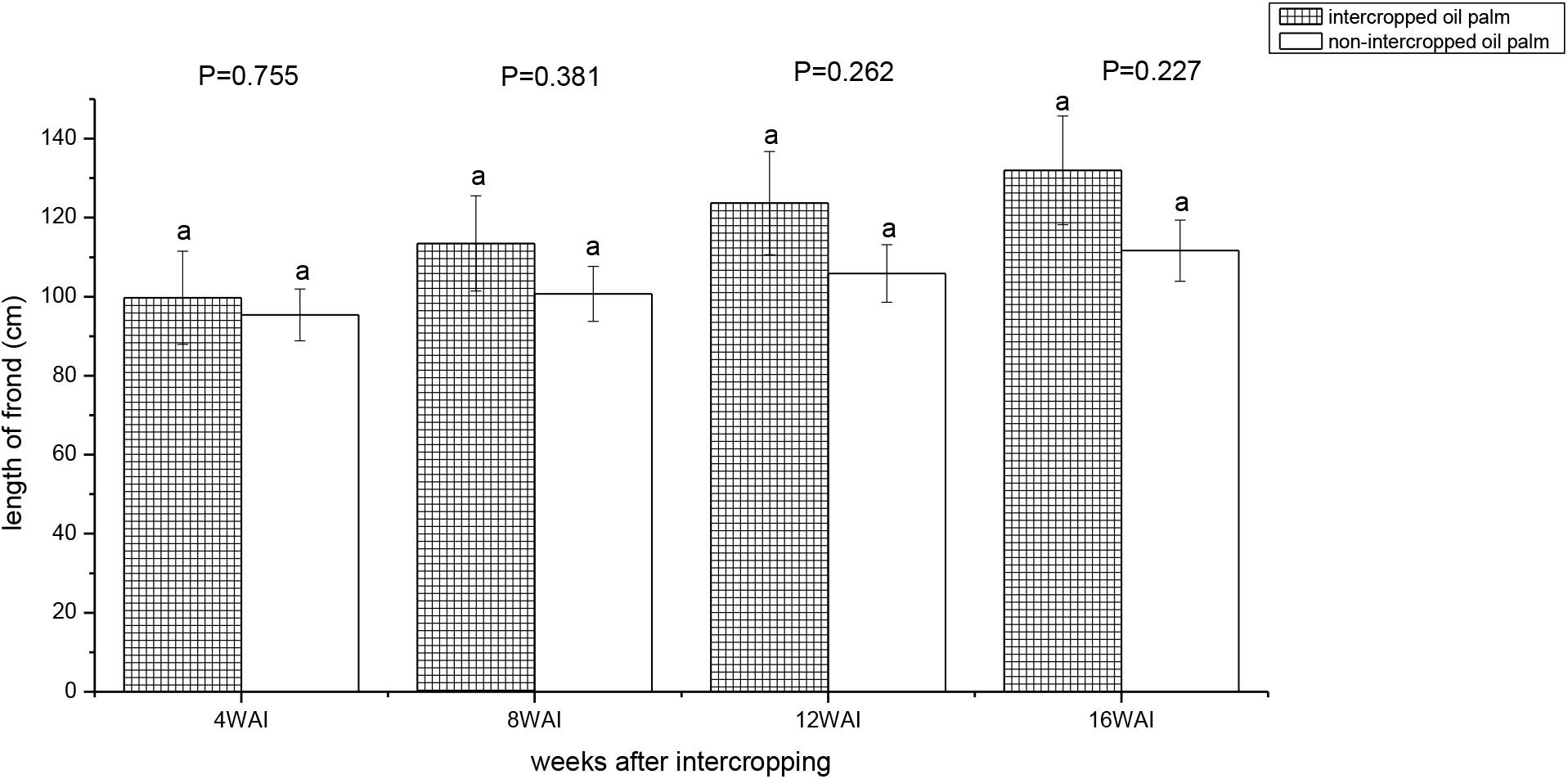
Length of frond of intercropped and sole juvenile oil palms at 4, 8, 12 and 16 weeks after intercropping. (n= 6) (2 years old).

**Fig. 4:**
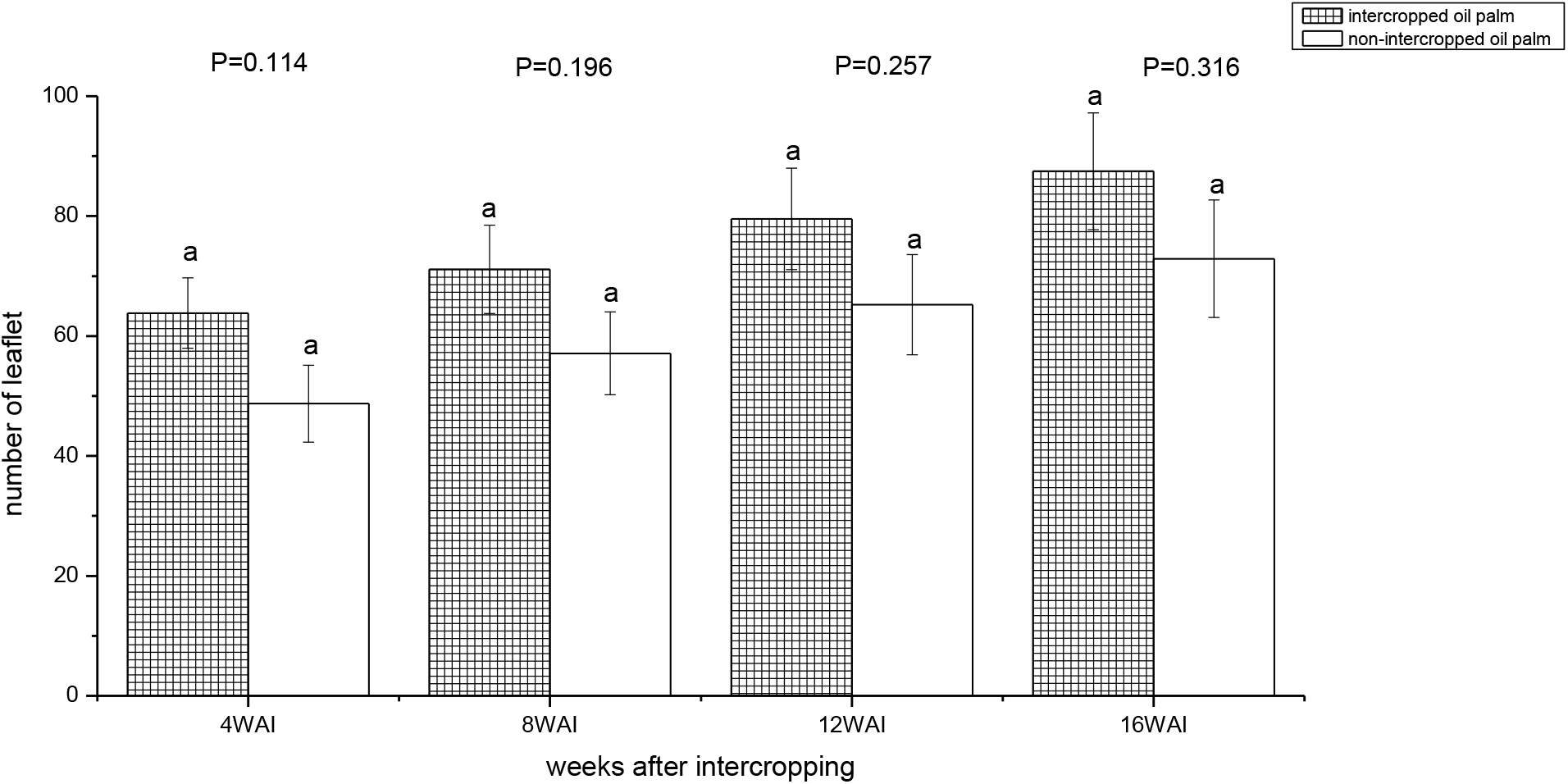
Number of leaflets of intercropped and sole juvenile oil palms at 4, 8, 12 and 16 weeks after intercropping. (n= 6) (2 years old).

**Fig. 5:**
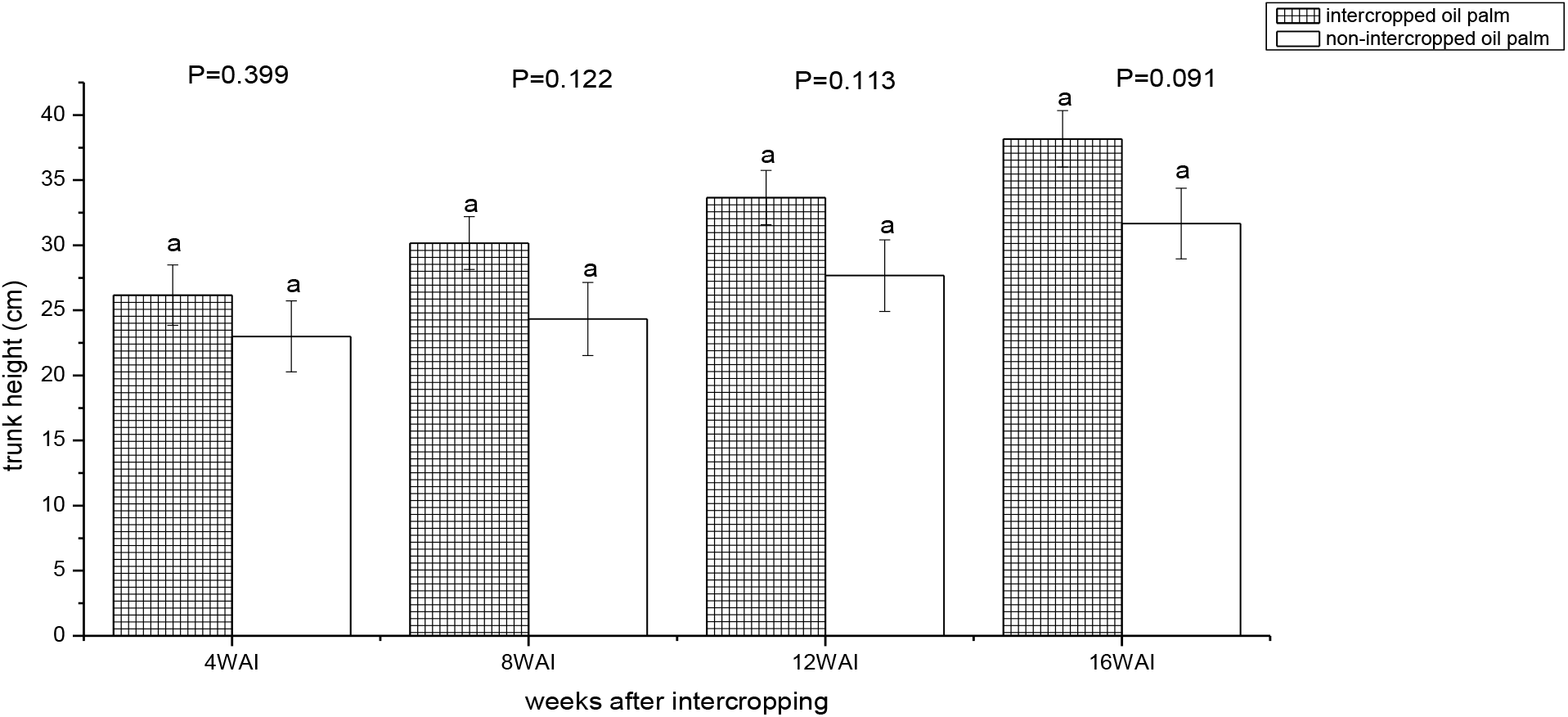
Trunk height of intercropped and sole juvenile oil palms at 4, 8, 12 and 16 weeks after intercropping. (n= 6).

**Fig. 6:**
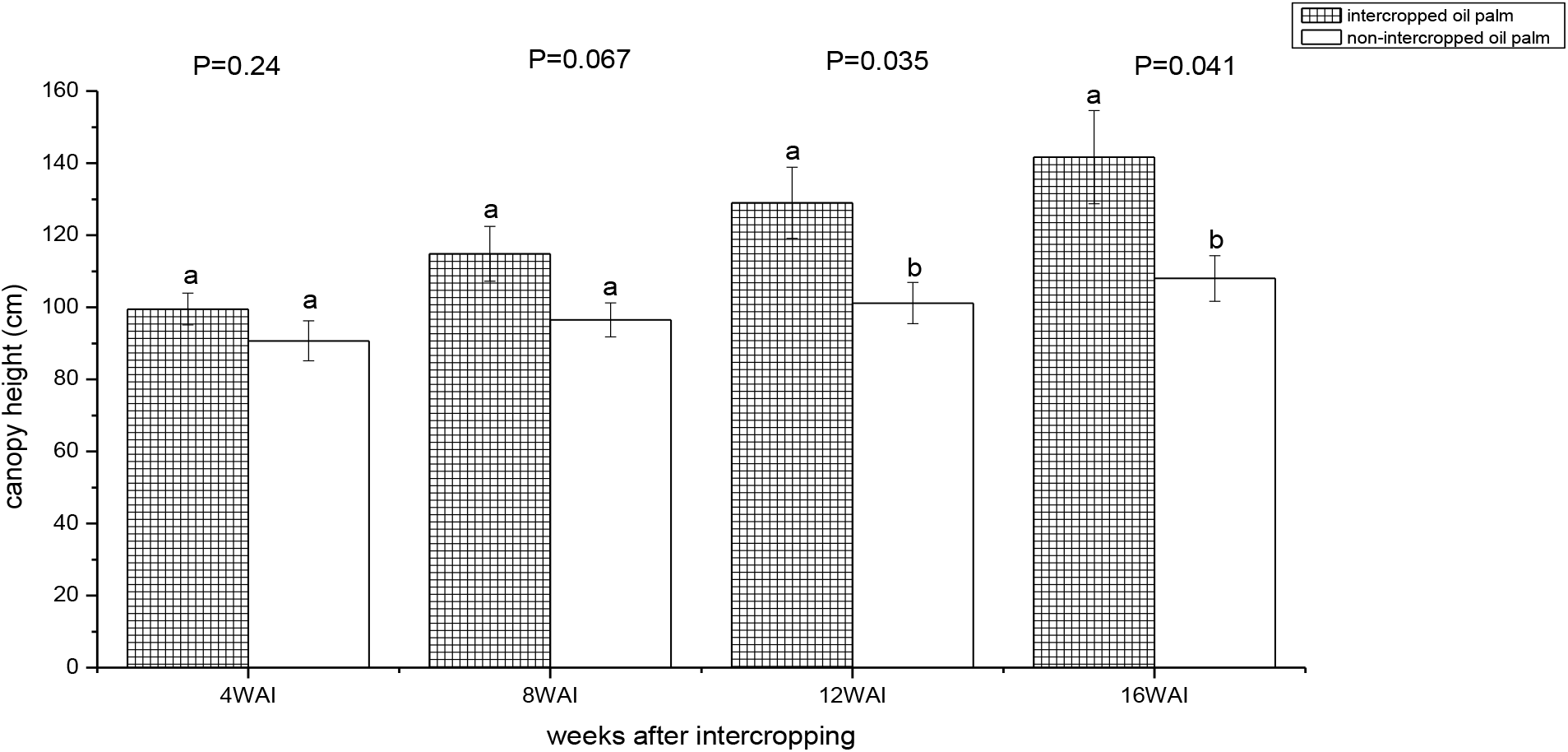
Canopy height of intercropped and sole juvenile oil palms after 4, 8, 12 and 16 weeks after intercropping. (n= 6) (2 years old).

Bars with the same letter are not significantly different at 95% degree of freedom (P≤0.05).

There was no significant difference between the canopy spread of the intercropped and sole juvenile oil palm at 4 weeks after intercropping through 16 weeks after intercropping, as none of the P-values are greater or equal to 0.05 as shown in Fig.1.

The number of fronds of intercropped juvenile oil palm was higher and statistically significant from that of the sole juvenile oil palm at 4, 8 and 12 weeks after intercropping. However, the mean numbers of frond were not statistically different at 16 weeks after intercropping (Fig. 2).

length of frond of intercropped and sole juvenile oil palms were not statistically different at 4 weeks after intercropping (WIA) through 16 WAI, with intercropped juvenile oil palm recording higher values (Fig. 3).

Fig. 4 shows that there was no statistically significant between the number of leaflets of intercropped and sole juvenile oil palm. However, the intercropped juvenile oil palm recorded higher mean number of leaflets at 4 WAI through 16 WAI.

The intercropped juvenile oil palm recording higher trunk height values; however, no significant difference was recorded at 4 WAI through 16 WAI as shown in Fig. 5.

The canopy heights of intercropped juvenile oil palms were higher than the sole juvenile oil palms; however, they were not statistically significant at 4 and 8 WAI. At 12 and 16 WAI, there were significant differences (Fig. 6). The results on the effects of intercropping fruit vegetables on the growth of juvenile oil palm during third year of plantation establishment are given in the Figure 7-12.

**Fig. 7:**
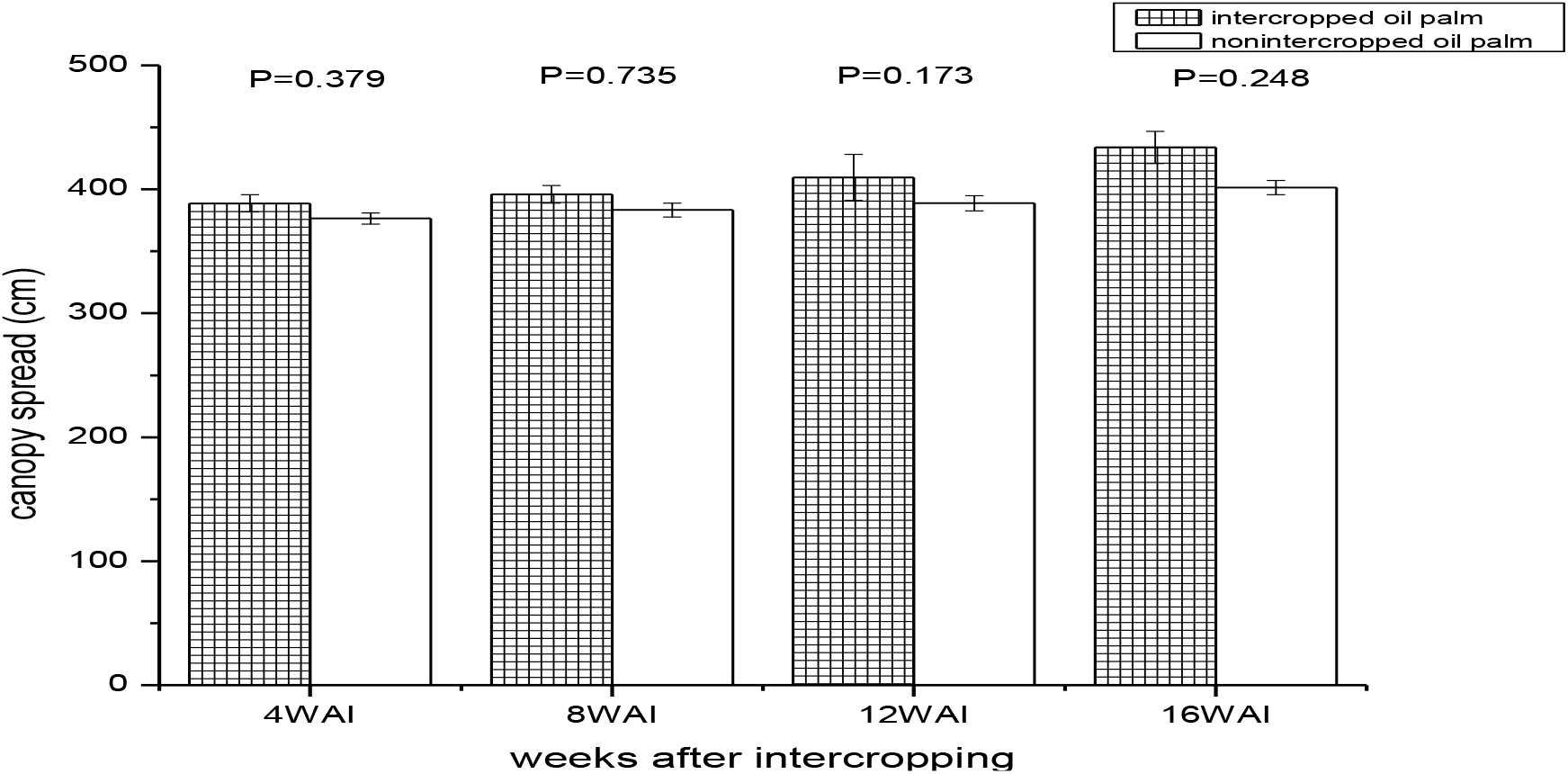
Effects of intercropping on canopy spread of intercropped and sole juvenile oil palms at 4, 8, 12 and 16 weeks after intercropping (n= 6) (2 years old).

**Fig. 8:**
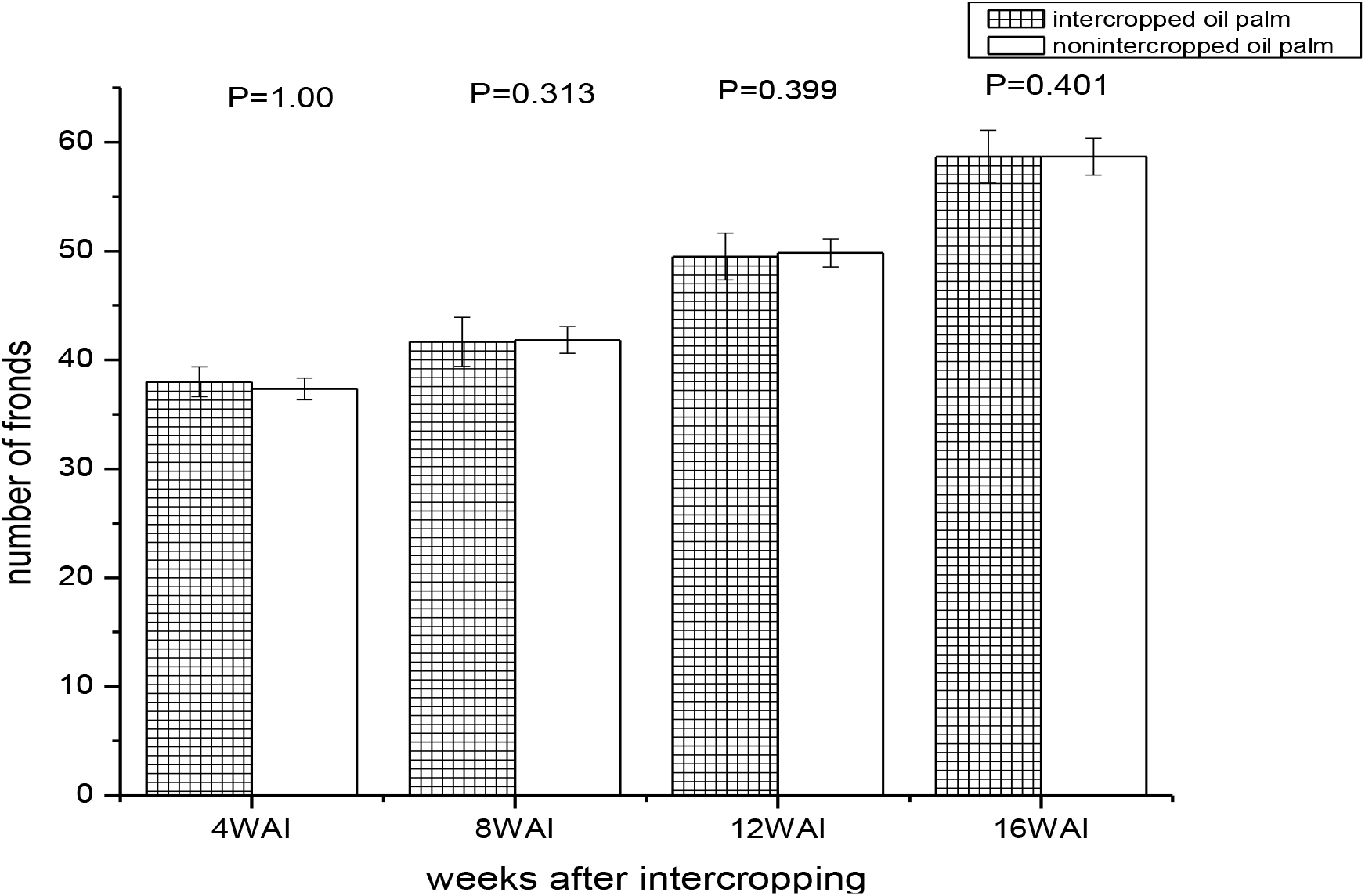
Effects of intercropping on number of fronds of intercropped and sole juvenile oil palms at 4, 8, 12 and 16 weeks after intercropping (n= 6) (2 years old).

**Fig. 9:**
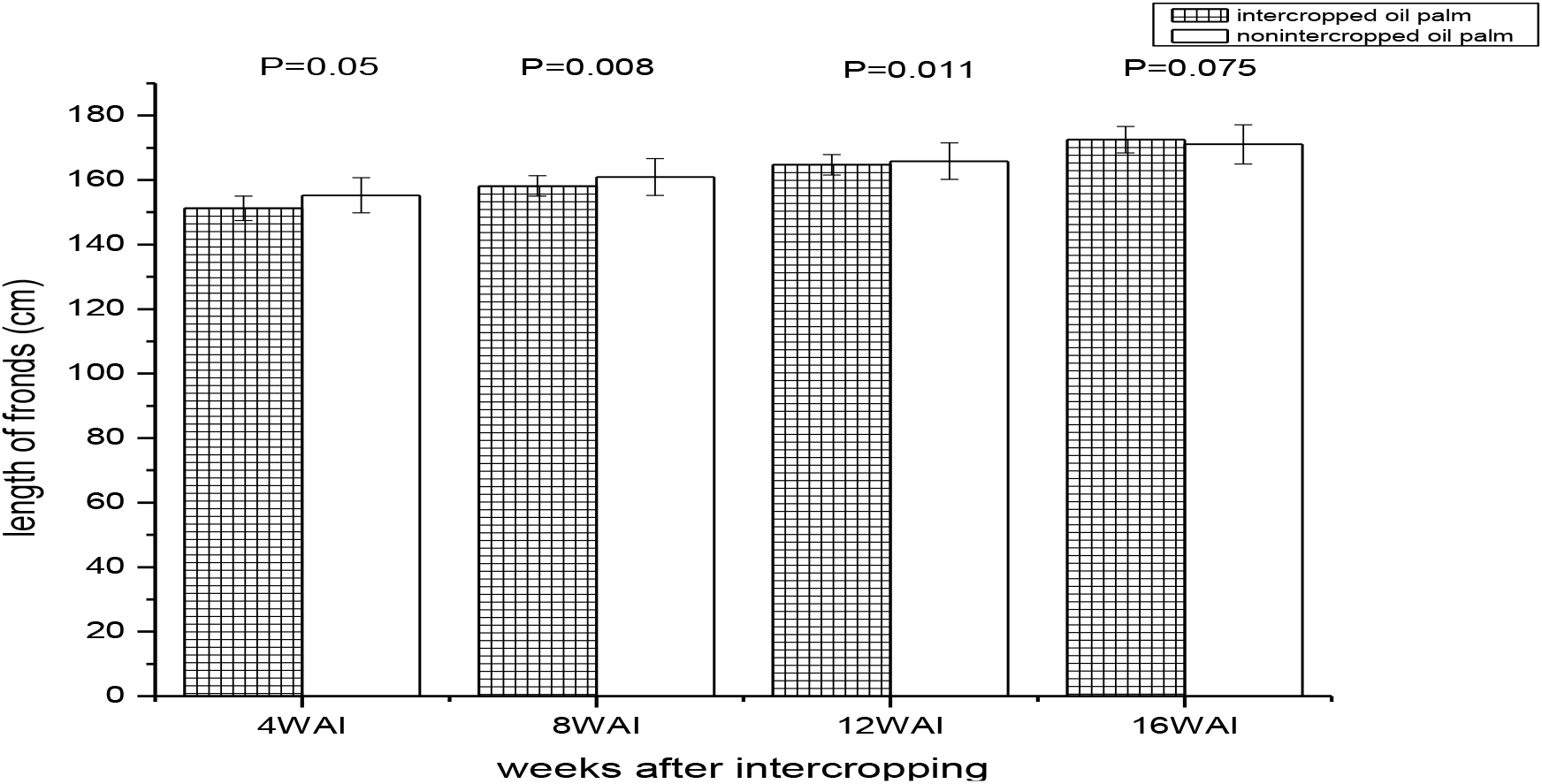
Effects of intercropping on length of fronds of intercropped and sole juvenile oil palms at 4, 8, 12 and 16 weeks after intercropping (n= 6) (2 years old).

**Fig. 10:**
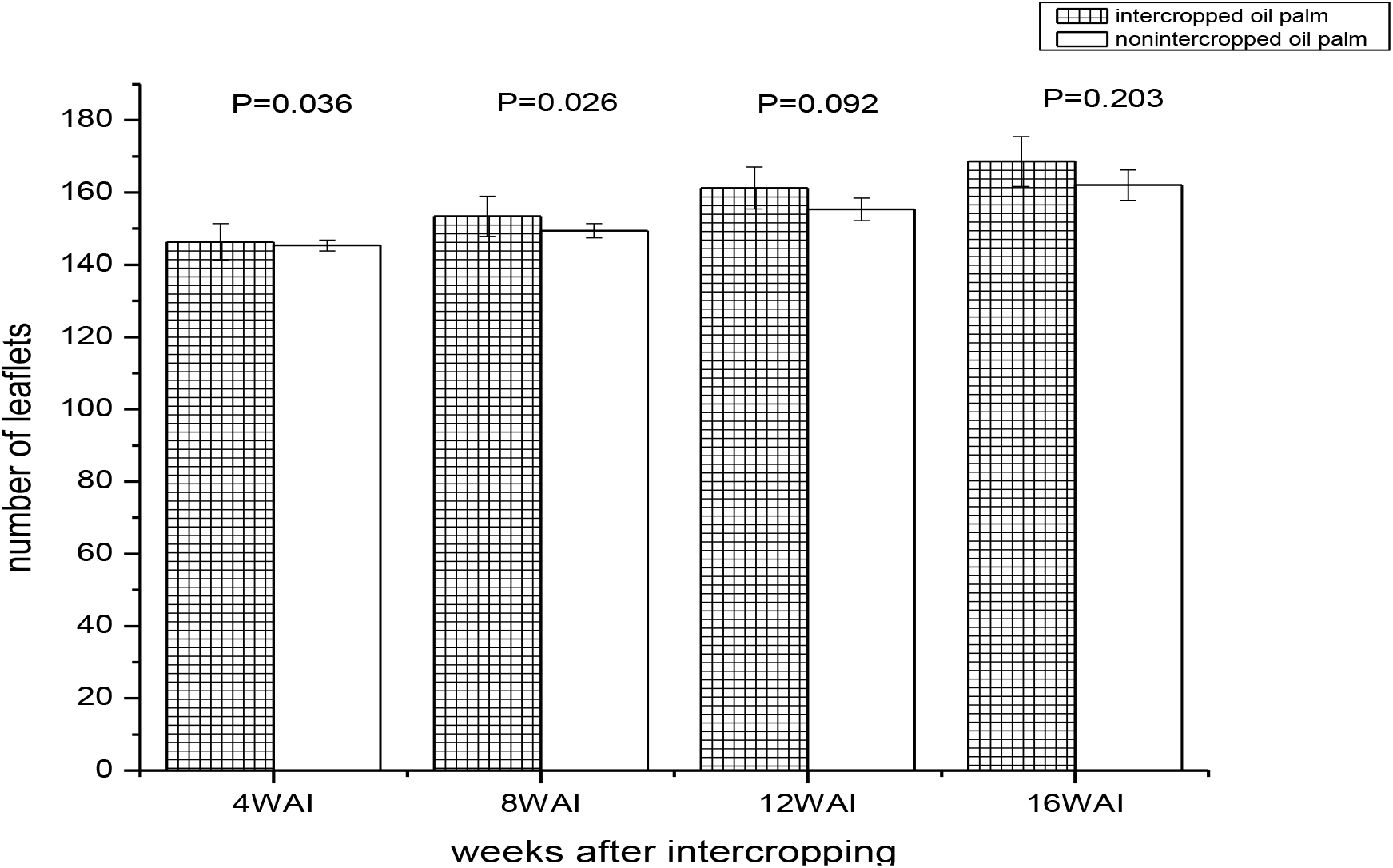
Effects of intercropping on number of leaflets of intercropped and sole juvenile oil palms at 4, 8, 12 and 16 weeks after intercropping (n= 6) (2 years old).

**Fig. 11:**
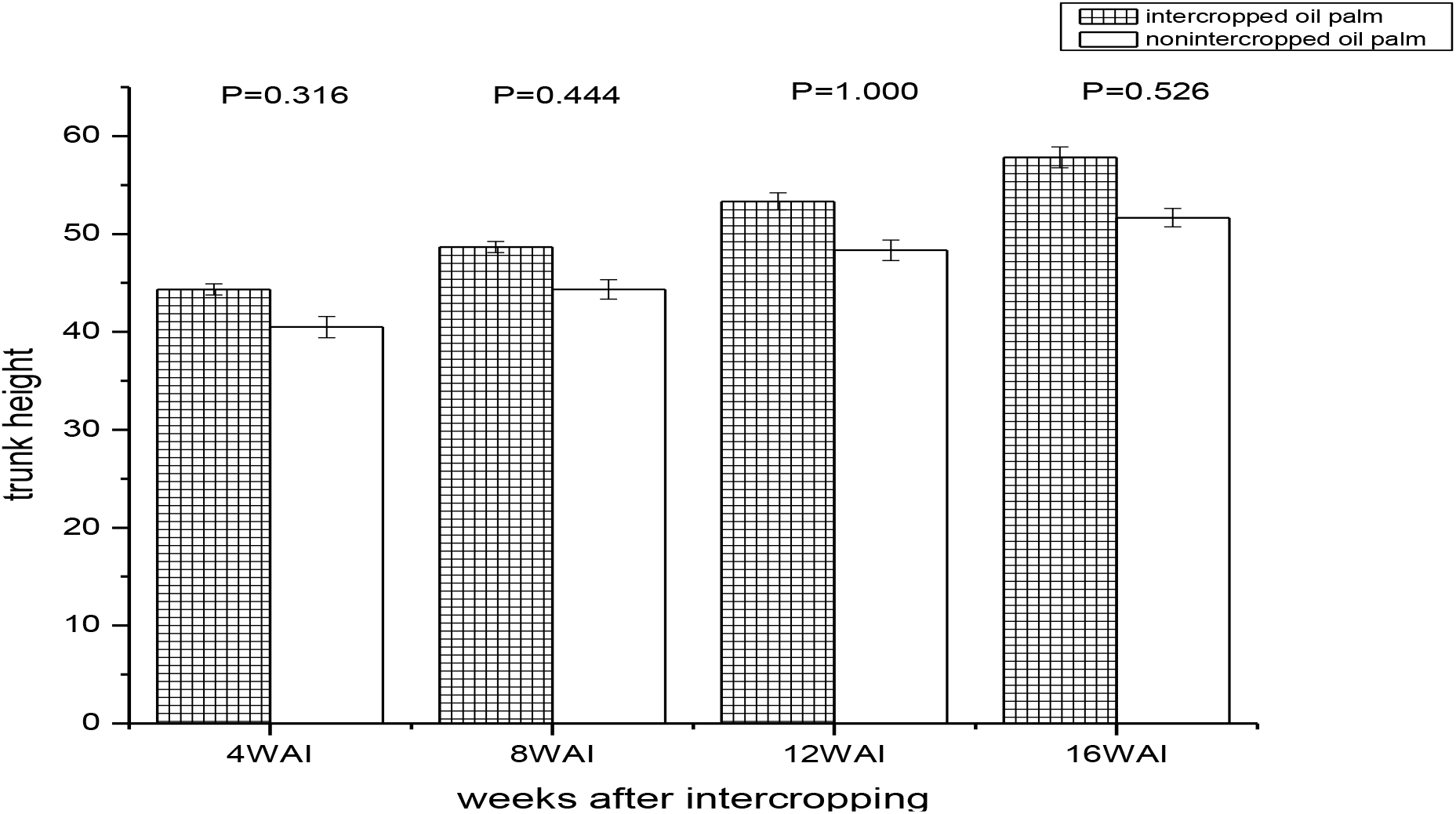
Effects of intercropping on trunk height of intercropped and sole juvenile oil palms at 4, 8, 12 and 16 weeks after intercropping (n= 6) (2 years old).

**Fig. 12:**
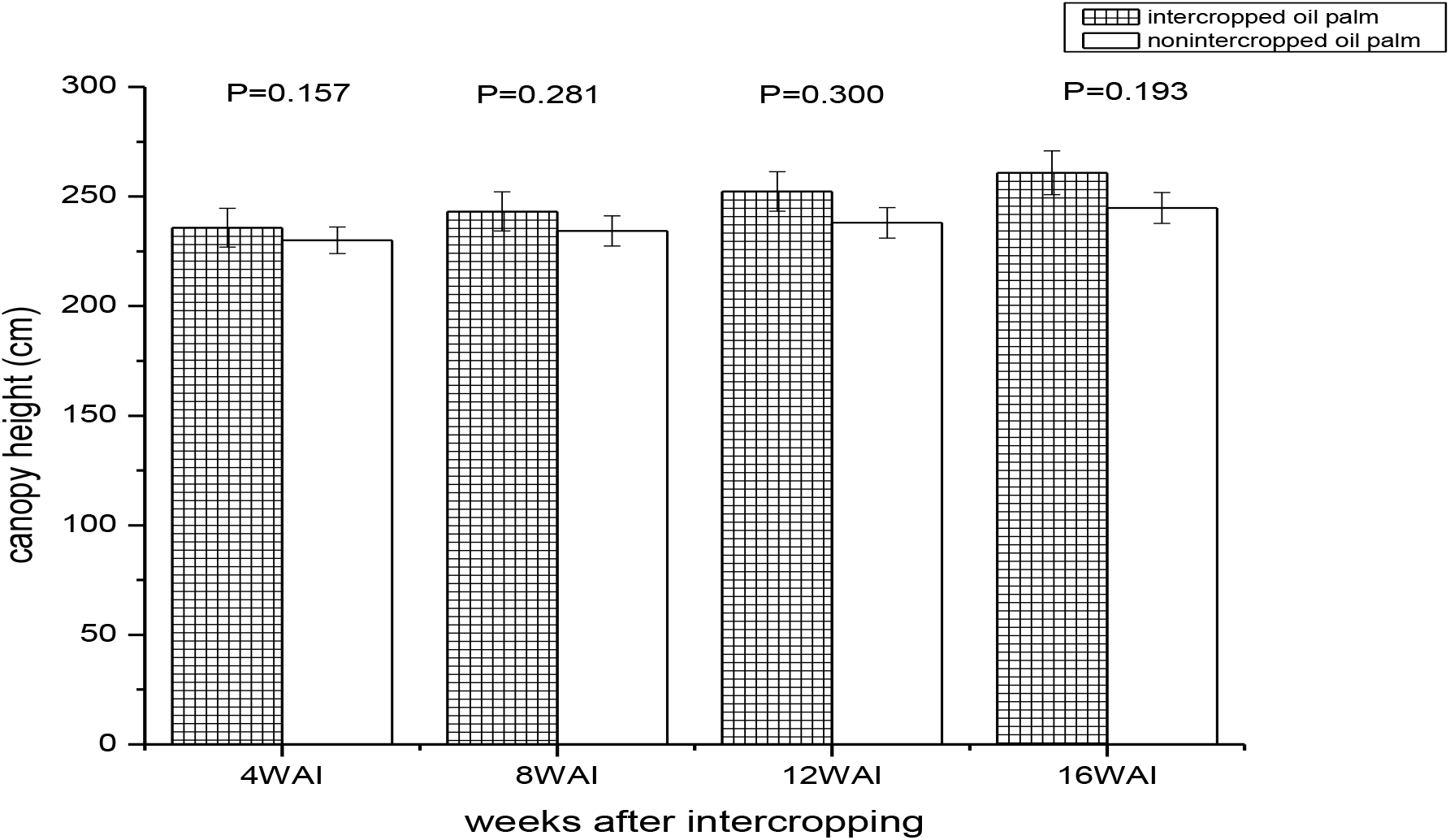
Effects of intercropping on canopy height of intercropped and sole juvenile oil palms at 4, 8, 12 and 16 weeks after intercropping (n= 6) (2 years old).

Bars with P≤0.05 are statistically significant at 95% degree of confidence.

The intercropped oil palms recorded higher canopy spread at 4, 8, 12 and 16 weeks after intercropping (WAI). However, there was no statistical significance between the canopy spread of intercropped and the sole juvenile oil palm from 4 WAI to 16 WAI as shown in Fig. 7.

At 4 WAI, the intercropped oil palm recorded higher number of fronds. The sole juvenile oil palms recorded higher number of fronds at 8 and 12 WAI. At 16WAI, the intercropped sole juvenile oil palm recorded the same mean number of fronds. The mean values of number of fronds for the intercropped and the non-intercropped juvenile oil palms were not statistically significant at 4 WAI through 16 WAI (Fig. 8).

At 4, 8 and 12 WAI, the sole juvenile oil palms recorded higher length of fronds and were statistically higher than the intercropped oil palms. At 16 WAI, the intercropped oil palms recorded higher length of fronds; however, it was not statistically significant (Fig. 9).

At 4, 8, 12 and 16 WAI, the intercropped juvenile oil palm recorded higher number of leaflets. However, they were statistically significant at 4 and 8 WAI as shown in Fig. 10.

The intercropped juvenile oil palms recorded higher trunk height at 4, 8, 12 and 16 WAI; however they were not statistically significant at 4 WAI through 16 WAI (Fig. 11).

Fig. 12 shows that intercropped juvenile oil palms recorded higher canopy spread at 4 WAI through 16 WAI; however, they were not statistically significant.

## Discussion

The results of the particle size indicated that all the experimental plots within the plantation were sand-loamy and by rating of FAO, the soil is medium textured. This quality indicated that the soil is appropriate for oil palm cultivation and this is in harmony with earlier work by Ukaegbu *et al.* (2015). Following the rating of Chude *et al.* (2011), not any of the experimental plots is deficient in nitrate before and after intercropping. The nitrate concentration ranged between 1.23 mg/l and 2.78 mg/l. the high concentration of nitrate is also associated with the high organic matter recorded in the experimental plots. The increase in available P observed soil of the intercropped juvenile oil palm plots was also reported by earlier work of Erhabor and Filson (2008) that available P level increased throughout the experimental periods in intercropped young oil palm plot, and the soil was rated to be low to medium in P level. There were moderate changes in the Ca and Mg, and there was no significant change in soil pH except in pepper (NGB 01312)-oil palm intercrop plot. Changes in soil organic matter and total organic matter fluctuated but did not follow definite trend. The intercropped juvenile oil palms recorded higher canopy spread, number of fronds, length of fronds, number of leaflets, trunk length and canopy height. However, at the end of 16 weeks after intercropping, there was no statistical difference in the mean values of the parameter examined except canopy height. This observation indicates that intercropping juvenile oil palm with fruit vegetables does not have detrimental effect on the oil palm; rather it helped to improve the growth. This better performance recorded in the intercropped juvenile oil palms may be attributed to more regular weeding carried out in the juvenile oil palm-vegetable intercrop plots. Furthermore, the initial minimum of 1 m space left fallow before planting the fruit vegetables reduced the competition between the juvenile oil palm and the fruit vegetables for nutrients in the soil and other growth resources. Similar result was obtained for the period of the second trial, there was no significant difference between the intercropped and sole juvenile oil palms latter at 16 weeks after intercropping. However, the sole juvenile oil palm recorded better performance in number and length of fronds than the intercropped ones probably because of no competition for resources by the vegetables intercropped.

Ogwuche *et al.* (2012) worked on the economies of intercropping natural rubber with arable crops as a panacea for poverty alleviation of rubber farmers. He reported that a higher annual increase in girth of rubber intercropped compared to the sole rubber plantation. He examined the influence of intercropping pattern on the girth of rubber plants and opined that at the end of every year during the intercrop, the girth of the rubber plants on both intercropped and non-intercropped plots increased; and was higher in the intercropped tree crops than the non-intercropped tree crops. This is in conformity with this study that intercropping of tree crop and arable crop such as vegetable crop would set in motion an increase in growth of the juvenile oil palm. They attributed this to an increase in organic matter content as a product of residues from the intercropped plants after harvest which promotes soil aeration and possible boost in soil nutrients required for the rubber growth and development. Decline in weed competition resulting from the intercrop may also be responsible.

More so, it could also be due to complementary of species interaction as opined by Esekhade (2003). He also indicated that intercropping tree crop with arable crops is profitable and can serve as a means of poverty alleviation among the rubber farmers if practiced. Similar report was also submitted by Giroh *et al.* (2010) who worked on efficiency and cost of production among gum Arabic farmers; that intercropping of food crops and tree crops is profitable. This report is in agreement with findings of this study that intercropping of vegetable crop and juvenile oil palms was profitable and simultaneously improved growth of the tree crop.

According to Oladokun (1990), among many tree crops that could be intercropped with food crops, most farmers prefer oil palm.

Udosen *et al*. (2006) investigated the performance of oil palm and different food crops combinations in four-year sequential intercropping in a rain forest/derived savanna transition zone of Nigeria. They observed that the different crop combinations intercropped with oil palm did not have any significant negative effect on the oil palm growth, number of bunches and fresh fruit bunch yield over the period. This observation is in agreement with findings of this study that intercropped juvenile oil palm did not show negative impact arising from the intercrop; rather it performed even better than the non-intercropped juvenile oil palm. They attributed this observation to the ability of the juvenile oil palm to maximize the available resources in the environment and did not have to compete for sunlight with the food crop which consequently resulted in good performance.

According to Okyere *et al.* (2014) who studied the lingering effect of intercropping on the yield and output of oil palm, reported that intercropping oil palm and food crop has no significant undesirable effect on the growth, development and yielding of the oil palm. He opined that intercropping does not absorb excessive nutrients from the field that would affect the nutrient requirements of the oil palms. He further attributed these findings to decomposition of crops residues after harvesting. More so, he suggested that the regular weeding of intercropped field and its eventual decomposition of weeds might have had added advantage to the nutrient availability for the growth of the oil palm even though that was not significant. Higher canopy height, number and length of frond, number of leaflets, trunk and canopy heights recorded in the intercropped than in the non-intercropped are in agreement with their observations as there was improvement in growth of the intercropped juvenile oil palms than the non-intercropped juvenile oil palm even though it was not significant.

This is also in accordance with an earlier work by Nuertey *et al.* (2000) who studied the economies of intercropping annual crops in oil palm plantation for small scale farmers that it is profitable to intercrop oil palm with food crops especially for the first two to four years when the oil palms are not fruiting as compared to sole cropping.

Famaye *et al*. (2012) investigated the effects of intercropping of coffee with rice and plantain at early stage of field establishment in Nigeria. They observed high growth performance in the intercrops and concluded that there were no harmful effects of intercropping of coffee with rice and food crops at early stage of field establishment. This is in agreement with findings of this study as intercropping of vegetable crops with juvenile oil palm has not shown negative effects on the growth of the intercropped and non-intercropped juvenile oil palm.

This finding is also corroborated by earlier works on the beneficial effect of intercropping some food crops with coffee (Okelana, 1982; Famaye, 2000 and 2005), cocoa, oil palm and kola (Adenikinju, 1980; Ofoli and Lucas, 1988; Okpala-Jose and Lucas, 1989; and CTA, 1993).

Famaye *et al.* (2012) reported that the yield obtained for food crops during intercrop of tree crops and food crops were as high as their sole crops. This further affirmed the beneficial impact on food crop production other than better morphological growth due to intercrop earlier pointed out as an advantage in intercropping (Herera and Harwood 1973; Okigbo and Greenland, 1975; Okigbo, 1977).

Putra *et al.* (2012) who probed the effect of intercropping style on growth of one-year-old oil palms reported that the food crops grown as intercrop with the juvenile oil palms did not inhibit the growth rate and performance of the juvenile oil palms which were the main crop. This is in concurrence with the findings of this study in which the intercropped juvenile oils were found to produce a better performance than the non-intercropped juvenile oil palms by recording higher canopy and trunk heights, canopy spread, and number and length of fronds. Similarly, this earlier report is also in harmony with the results of this study that though the intercropped juvenile oil palms had better growth performances than the non-intercropped palm during the second- and third-year of field establishment, the differences were largely not significant.

Contrary to the findings of this study, Rafflegeau *et al.* (2010) reported negative impact of annual crop intercrop with immature oil palms. They pointed out that the presence of food crops on oil palm plantations at the immature stage resulted in nitrogen and potassium deficiencies which persisted even when the plantation reaches the production stage especially without appropriate annual fertilization.

## Conclusion

Intercropping of fruit vegetables at a minimum of 1 m away from the row juvenile oil palms did not result in deleterious effects in the young oil palms; rather, it encouraged their growth and development.

This study established that fruit vegetables namely tomato, pepper, okra, and eggplant can successfully be intercropped with juvenile oil palm intercrops during the second and third year of field establishment of the oil palm. This success in intercrop was achieved by planting the fruit vegetables at a minimum of 2 m away from the juvenile oil palm.

